# Intrinsic spectrally-dependent background in spectroscopic visible-light optical coherence tomography

**DOI:** 10.1101/2020.09.13.294876

**Authors:** Ian Rubinoff, Roman V. Kuranov, Hao F. Zhang

## Abstract

Visible-light optical coherence tomography (vis-OCT) enabled new spectroscopic applications, such as retinal oximetry, as a result of increased optical absorption and scattering contacts in biological tissue and improved axial resolution. Besides extracting tissue properties from back-scattered light, spectroscopic analyses must consider spectral alterations induced by image reconstruction itself. We investigated an intrinsic spectral bias in the background noise floor, which is hereby referred to as the spectrally-dependent background (SDBG). We developed an analytical model to predict the SDBG-induced bias and validated this model using numerically simulated and experimentally acquired data. We found that SDBG systemically altered the measured spectra of blood in human retinal vessels in vis-OCT, as compared to literature data. We provided solutions to quantify and compensate for SDBG in retinal oximetry. This work is particularly significant for clinical applications of vis-OCT.

## 1. Introduction

Optical coherence tomography (OCT) [1] detects back-scattered light to image biological tissue at microscopic resolutions noninvasively. In spectral-domain OCT (SD-OCT), three-dimensional (3D) images are reconstructed using the Discrete Fourier transform (DFT) of a sampled interference spectrum. Visible-light OCT (vis-OCT) [2] is a rapidly evolving SD-OCT technology that operates within the visible-light wavelength range to increase axial resolution and spectroscopic tissue contrast, as compared with near-infrared OCT (NIR-OCT). These benefits enabled new spectroscopic vis-OCT applications, including oximetry [3, 4], detection of tissue ultrastructure [5], and investigating various neuropathologies [6, 7]. Spectroscopic vis-OCT computes a series of short-time Fourier Transforms (STFT) using moving spectral windows with reduced bandwidth across the entire spectral interference fringes. Performing STFT enables the reconstruction of a series of sub-band spectral images with a reduced axial resolution.

To accurately extract tissue spectral information, it is essential to eliminate influence from the OCT image reconstruction itself. In most SD-OCTs, a grating-based spectrometer samples the interference fringe almost linearly in the wavelength (λ) space, which is inversely proportional to the wavenumber (*k*) space. The optimal axial resolution requires the interference fringe to be interpolated linearly in *k* space [8, 9]. We found that linear-in-*k* interpolation generates spectrally-dependent background (SDBG), an intrinsic bias of the vis-OCT background noise floor that can alter spectroscopic measurements.

The prevalence of linear-in-*k* interpolation in SD-OCT makes the systemic nature of the SDBG highly significant towards the spectroscopic OCT research community. Various background biases have been previously reported [10–14]. For example, in polarization-sensitive OCT (PS-OCT) the noise floor varied in different polarization channels and, therefore, comparing the polarization channels required noise floor correction [11]. Although researchers previously recognized the existence of SDBG and applied empirical corrections [12, 14], a thorough investigation of SDBG’s origin, influence on spectroscopic vis-OCT, and correction techniques have yet to be conducted.

In this work, we first theoretically derive the systemic bias of SDBG caused by linear-in-*k* interpolation of the interference fringe. We simulate SDBG in our vis-OCT system and compare it with experimentally acquired SDBG. Then, we establish a minimum fringe upsampling rate in spectroscopic vis-OCT that removes many depth-dependences of the SDBG, increasing simplicity of SDBG correction. Finally, we investigate the influence of SDBG on vis-OCT retinal oximetry and apply an SDBG correction strategy. This work establishes important principles and consequences of widely-used data processing in spectroscopic vis-OCT and all other SD-OCTs, informing a broad range of biophotonic applications.

## 2. Origin and derivation of SDBG

### 2.1 Wavenumber dispersion in spectrometer detection

In SD-OCT, the noise-free interference fringe (neglecting the DC component) can be written as

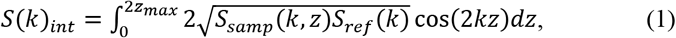

where *z* is the depth of the collected back-scattered photons from the zero-delay; *z*_*max*_ is the maximum imaging depth [15]; *S*_*samp*_(*k*, *z*) is the power spectrum of the back-scattered light from the depth *z*; and *S*_*ref*_(*k*) is the power spectrum of the reference arm. *S*(*k*)_*int*_ is spatially dispersed onto a one-dimensional (1D) pixel array in the spectrometer as a function of *k*(*x*) across the range from *k*_*start*_ to *k*_*end*_, which is determined by the grating equation and spectrometer optics [16]. Here, *x* is the spatial coordinate along the 1D array ranging from 0 to *N*Δ*x*, where *N* is the the total number of pixels and Δ*x* is the width of each pixel.

The spatial dispersion of the spectrometer is represented by 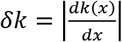, which we refer to as the *k* spacing. It is also important to consider a hypothetical uniform dispersion across the same spectral range, 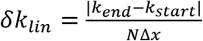, which is constant. The dimensionless ratio, 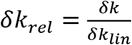, referred to as relative *k* spacing, describes how the spectrometer deviates from ideal uniform *k* dispersion across each pixel. Indeed, a larger *δk*_*rel*_ indicates more *k*-space bandwidth per unit pixel Δ*x*, while a smaller *δk*_*rel*_ indicates less *k*-space bandwidth per unit pixel. The interference fringe in the Eqn. 1 sampled by the 1D pixel array (without considering spectrometer roll-off [17]) can be written as

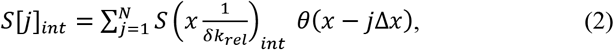

where *j* = 1, 2, … *N* is the array pixel index, and *θ*(*x* − *j*Δ*x*) is the Dirac comb function with a period Δ*x*. The spectrometer pixel array samples with a uniform period in *x*, but the fringe is a function of 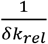. Such a non-uniform sampling of the *k* space results in a phase nonlinearity in *S*[*j*]_*int*_ [18], which distorts *S*[*j*]_*int*_ and reduces image axial resolution. For optimal image reconstruction, the phase of *S*[*j*]_*int*_ must be made linear-in-*k*, which we will discuss in Section 2.2.

Quantifying 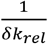 for a spectrometer is valuable for understanding the distortion of *S*[*j*]_*int*_. In this work, we measured 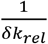 digitally sampled by the spectrometer: 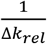. From here on, *δk*_*rel*_ refers to the *k* spacing in the continuous domain, while Δ*k*_*rel*_ refers to the *k* spacing in the discrete domain. Briefly, we found the *k* distribution on the pixel array *k*[*j*]_*map*_ using spectral calibration lamps [17, 18]. Next, we obtained the sampled *δk*, defined as Δ*k*, by calculating the absolute change in *k*[*j*]_*map*_ with *j*. Hence, the sampled *δk*_*lin*_ is

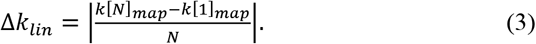

From this information, we obtained the sampled *δk*_*rel*_, defined as 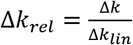.

In Figure 1a, we plot measured 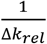 and 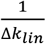 as a function of pixel *j* for a commercial vis-OCT spectrometer (Blizzard SR, Opticent Health, Evanston, IL). Briefly, the spectrometer collimates light from a single-mode fiber output. The collimated light is diffracted by a transmission grating. The diffracted light is focused on a 2048-pixel camera (OctoPlus, Teledyne E2V, UK) placed at the focal plane of the diffracted light. The spectral detection range is from 506 nm to 613 nm. We measured the full spectrum roll-off as −4.8 dB/mm and confirmed that aberrations at the focal plane were minimized by measuring the spectrally-dependent roll-off (SDR). [17].

**Fig. 1.**
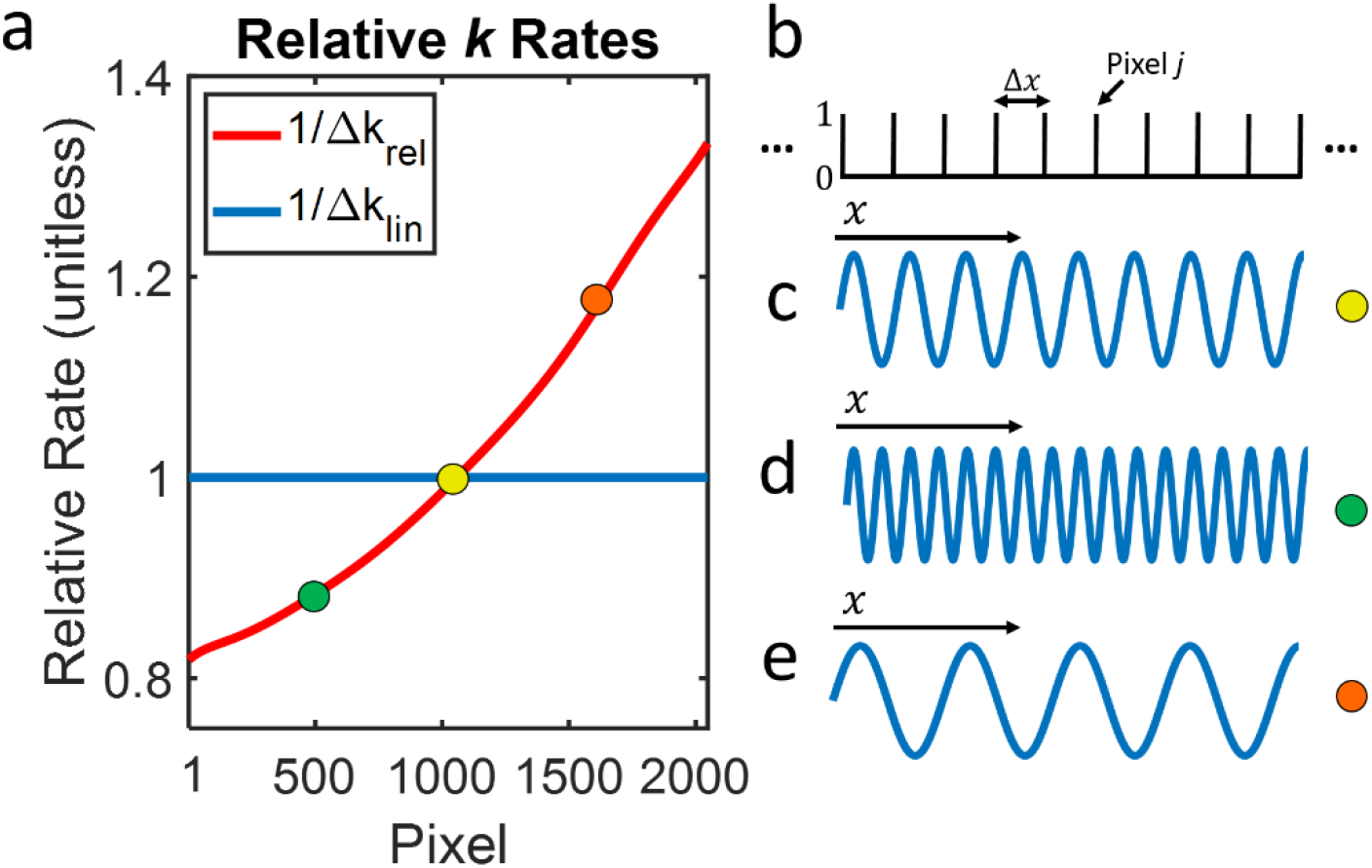
Influence of grating dispersion on interference fringe (a) Digitized relative *k* rates of a spectrometer. Colored dots represent locations of pixel segments in (c-e); (b) illustration of Dirac comb sampling by pixel array with period Δ*x*; (c) Interference fringe plotted at the location of the yellow dot in panel a; (d) Interference fringe plotted at the location of the green dot in panel a; (e) Interference fringe plotted at the location of the orange dot in panel a.

From Fig. 1a, is evident that 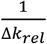 is smaller for shorter wavelengths and larger for longer wavelengths. Physically, this can be considered as a compression of the *k*-space for shorter wavelengths and an expansion of the *k*-space for longer wavelengths, as shown in Figs. 1b-1e. Fig. 1b shows a segment of the Dirac Comb from Eqn. 2, representing sampling by 10 pixels with a period of Δ*x*. Fig. 1c shows a single sine wave, representing a segment of 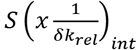, where 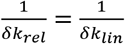 (highlighted by the yellow dot in Fig. 1a). All sine waves are plotted along the same *x*-axis from Fig. 1b. Fig. 1d shows the same 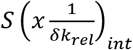 as in Fig. 1c, but for a segment where 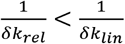 (highlighted by the green dot in Fig. 1a). This condition causes a compression of the *k*-space. Therefore, more cycles of 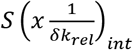 are observed comparing with Fig. 1c. Since the pixel array samples with uniform period Δ*x*, which is independent of *δk*_*rel*_, dispersing more *k*-space per pixel comes at the expense of acquiring fewer *k*-space samples. As shown in Eqn. 2, 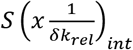 experiences a relatively sparse discrete sampling rate when 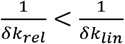. In this way, we show that the *k*-space sampling rate is proportional to 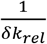 (and estimated by 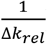). Fig. 1e shows 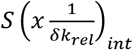 for a segment where 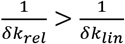 (highlighted by the orange dot in Fig. 1a), where expanded *k*-space results in a relatively dense *k*-space sampling rate when 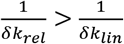. Other than the spectral distortions described above, uniform sampling by the pixel array itself does not introduce any new distortions (not considering aliasing and roll-off).

### 2.2 Linear-k interpolation and resampling

A linear-in-*k* version of *S*[*j*]_*int*_ can be estimated by digitally redistributing the locations of *k*-space samples. Specifically, *S*[*j*]_*int*_ is interpolated to linear-in-*k* by calculating unknown values of *S*[*j*]_*int*_ from an estimated continuous version of itself. Due to its simplicity and computational efficiency, a popular SD-OCT fringe interpolation is linear interpolation (LI) [19], which is the focus of our analysis. However, the mathematical principles and fundamental conclusions derived here remain valid for other interpolations, such as cubic spline. A discrete LI consists of three steps: up-sampling, low-pass filtering, and down-sampling [20].

As shown above, *S*[*j*]_*int*_ is initially down-sampled at shorter wavelengths and up-sampled at longer wavelengths according to 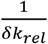. Phase distortion of *S*[*j*]_*int*_ can be removed by an inverse operation. Therefore, *S*[*j*]_*int*_ is resampled at a rate inverse to 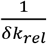, which is in the up-sampling and down-sampling steps in LI. As illustrated in Fig. 1, LI results in an expansion of shorter wavelengths and compression of longer wavelengths exactly inverse to their original distortions. This can be described mathematically as

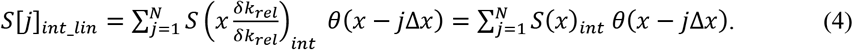

From Eqn. 5, the noise-free interference fringe is resampled linearly in *k* and *x*, canceling sampling-based distortions. Finally, the low-pass filtering step in LI uses a triangle function

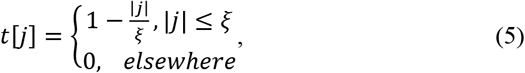

where *ξ* is the order of interpolation [21]. We show the influence of Eqn. 5 in Section 3.3.

### 2.3. Noise in SD-OCT

In SD-OCT, the additive noise consists of shot noise, dark noise, readout noise, and relative intensity noise (RIN) and is assumed to be Gaussian distributed at each pixel [22]. Unlike sampled fringe *S*[*j*]_*int*_, the additive noise is not correlated with *k*. Therefore, we modify representation of the signal detected by the spectrometer to

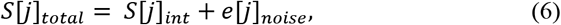

where *e*[*j*]_*noise*_ is the additive noise in SD-OCT.

We confirmed the Gaussian noise distribution by acquiring images using our vis-OCT system (40 *μs* exposure time, 5000 A-lines, 0.15 *μW* laser power from reference arm) without any sample in the sample arm. After normalizing the detected noise by the spectral shape of the light source (NKT Photonics, SuperK 150 MHz), we found that our measured noise followed the Gaussian distribution with a mean near 0 and a standard deviation (*σ*) of 0.79 [arb. Units]. Additionally, it has been shown in SD-OCT that a supercontinuum laser source contributes pink RIN noise [22]. In our experimental measurements in Section 3, we demonstrate that although pink noises exist, SDBG bias and the general spectral profile of the background is dominated by the white noise component.

### 2.4. Interpolation of noise in SD-OCT

To understand how interpolation leads to SDBG, we must consider a complete representation of the SD-OCT signal

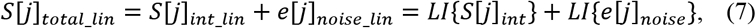

where *S*[*j*]_*total*_*lin*_ is the linearly interpolated signal detected by the spectrometer; *S*[*j*]_*int*_*lin*_ is the linearly interpolated interference fringe; *e*[*j*]_*noise*_*lin*_ is the linearly interpolated noise; and *LI* is the linear interpolation operator. *S*[*j*]_*int*_ and *e*[*j*]_*noise*_ are interpolated independently.

A useful property of white noise is that its autocorrelation is proportional to the Dirac delta function. First, we denote the autocorrelation of *e*[*j*]_*noise*_ as 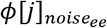. To investigate 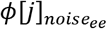 in the same way as the sampled interference fringe, we write 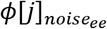 as

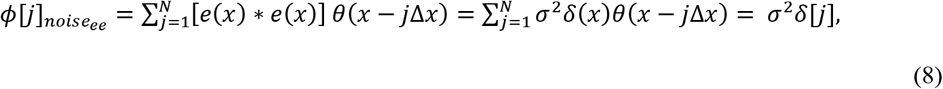

where *e*(*x*)_*noise*_ is the SD-OCT noise as a function of continuous space *x*; * is the continuous convolution operator; *σ*^2^ is the variance of the noise; and *δ*(*x*) and *δ*[*j*] are the continuous and discrete Dirac delta functions, respectively. Eqn. 8 is valid because *e*[*j*]_*noise*_ is a wide-sense stationary signal [20]. Unlike the interference fringe, neither *e*[*j*]_*noise*_ nor *e*(*x*) are correlated with *k*. As such, from the perspective of the spectrometer array, *e*(*x*), *e*[*j*]_*noise*_, and 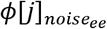 are all assumed linear with the pixel index. However, interpolating *S*[*j*]_*int*_ still necessitates linear-in-*k* interpolation of *e*[*j*]_*noise*_. Similar to Eqn. 4, we can write the interpolated noise as

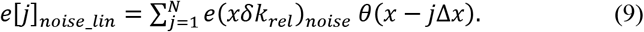

Now, we can write the autocorrelation of the interpolated noise, as

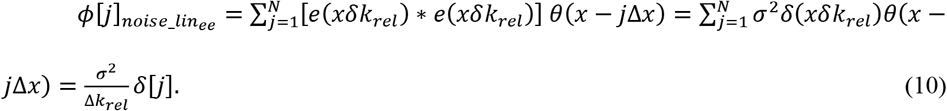

Indeed, expansion or compression by *δk*_*lin*_ does not linearize *k* domain sampling for *e*[*j*]_*noise*_ like it does for *S*[*j*]_*int*_. Eqn. 11 shows that the amplitude of 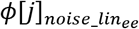 is determined by *δk*_*rel*_ and scaled by 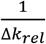.

### 2.5. SDBG in the reconstructed depth spectrum

Since we are interested in the depth-resolved spectral signatures of vis-OCT signals, we need to investigate the STFT of *S*[*j*]_*total*_*lin*_ as

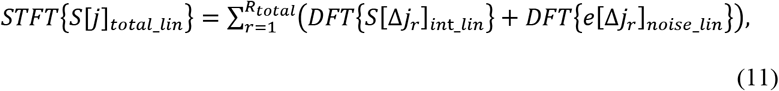

where *r* is the STFT sub-band spectral window and Δ*j*_*r*_ are the sample indexes under the full-width-half-maximum (FWHM) bandwidth of spectral sub-band window *r*. *S*[Δ*j*_*r*_]_*int*_*lin*_ and *e*[Δ*j*_*r*_]_*noise*_*lin*_ are the windowed versions of *S*[*j*]_*int*_*lin*_ and *e*[*j*]_*noise*_*lin*_, respectively. We can compute STFT of 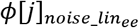 as

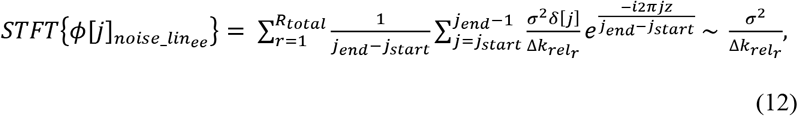

where *z* is the transformed index of *j* representing the depth; *i* is the complex number; *j*_*start*_ and *j*_*end*_ are the first and last indexes of Δ*j*_*r*_, respectively; and 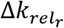 is the windowed version of Δ*k*_*rel*_. Applying the Wiener-Khinchen theorem, the DFT of 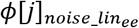 for each sub-band is the power spectral density (PSD) of *e*[*j*]_*noise*_*lin*_ under each sub-band, referred to as 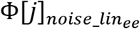 [20]. Since the result of Eqn. 12 is a positive constant for each *r*, we can relate the PSD to its amplitude spectrum (spectrally dependent A-line or SDA-line: *A*[*z*, Δ*j*_*r*_]_*noise*_*lin*_) using

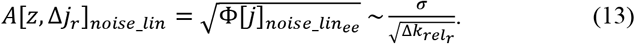

Then Eqn. 11 can be rewritten as

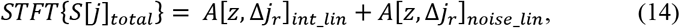

where *A*[*z*, Δ*j*_*r*_]_*int*_*lin*_ is an SDA-line reconstructed from the fringe. SDBG is therefore a bias of the noise floor proportional to 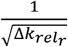.

## 3. Results

### 3.1. Simulated SDBG

We simulated SDBG for the Blizzard SR spectrometer by generating a Gaussian white noise (*e*[*j*]_*noise*_) and processing it following an STFT spectroscopic vis-OCT reconstruction procedure [2]. Briefly, we generated *e*[*j*]_*noise*_ with *N* = 2048 pixels and an SD of 0.79 (arb. units, same as measured in Section 2.3). Then, we up-sampled *e*[*j*]_*noise*_ six times using FT zero-padding [19] and linearly resampled to obtain *e*[*j*]_*noise*_*lin*_. Finally, we applied STFTs using *R*_*total*_ = 24 Gaussian windows spaced equidistantly in the *k* space from 523 nm to 591 nm. Each sub-band had the same FWHM bandwidth, corresponding to a 13-nm FWHM bandwidth for a sub-band centered at 556 nm. We repeated this processing 5000 times with and without the LI step and averaged all the respective 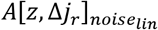.

Fig. 2a shows the simulated *A*[*z*, Δ*j*_*r*_]_*noise*_*lin*_ with LI after normalizing by its average spectral amplitude 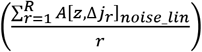 and average depth amplitude between z = 500 μm and z = 800 μm 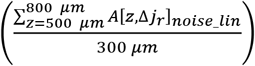. From here on, all *A*[*z*, Δ*j*_*r*_]_*noise*_*lin*_ are plotted after this normalization. The simulated *A*[*z*, Δ*j*_*r*_]_*noise*_*lin*_ has 24 background values, each representing the *r*^*th*^ STFT sub-band. We color-code the central wavelength of each sub-band, as shown by the color bar. Although all *A*[*z*, Δ*j*_*r*_]_*noise*_*lin*_ are approximately constant with z, the mean amplitude from shorter (green) wavelengths are higher than those from longer (orange) wavelengths. We visualize the simulated SDBG bias in Fig. 2b (red dashed line), which we approximate as the depth-averaged *A*[*z*, Δ*j*_*r*_]_*noise*_*lin*_ between 500 μm and 800 μm:

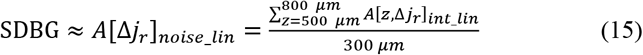

**Fig. 2.**
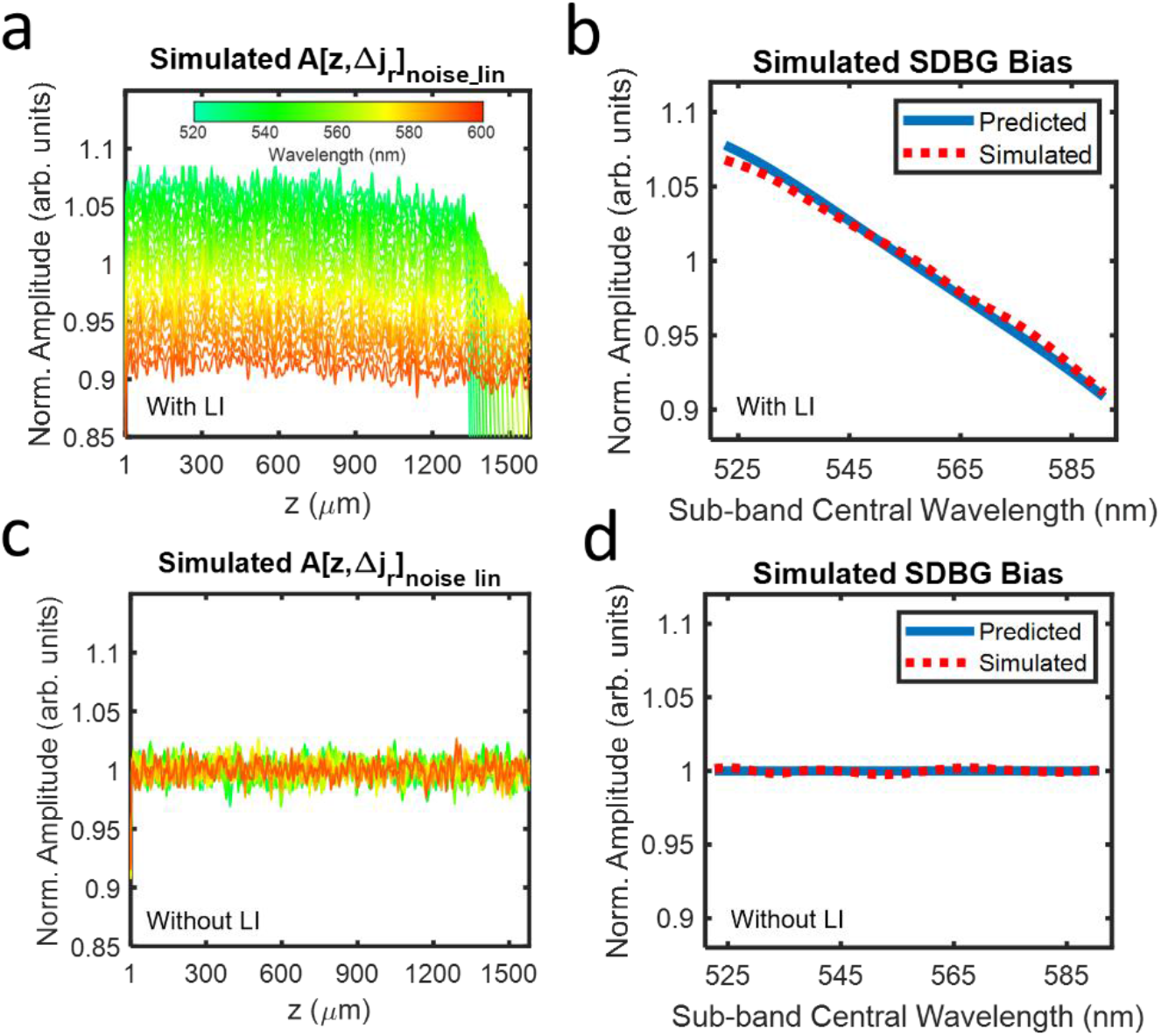
Simulated SDBGs in vis-OCT. (a) Simulated background SDA-lines with LI; (b) predicted SDBG bias (blue line) and simulated SDBG bias (red dashed line) with LI from z = 500 μm to 800 μm; (c) simulated background SDA-lines without LI; (d) predicted SDBG bias (blue line) and simulated SDBG bias (red dashed line) without LI from z = 500 μm to 800 μm

The blue line in Fig. 2b shows the SDBG bias predicted by the spectrometer’s 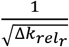, which agrees well with the simulated *A*[Δ*j*_*r*_]_*noise*_*lin*_. Fig. 2b and all plotted *A*[Δ*j*_*r*_]_*noise*_*lin*_ are normalized by their average spectral amplitude, as done in Fig. 2a. We noted that there is a small difference between the SDBG biases at the shortest wavelengths, although this difference is < 1% of the predicted value. This difference may be caused by *σ* not being perfectly constant with sub-band center wavelength or a minute depth-dependence of *A*[*z*, Δ*j*_*r*_]_*noise*_*lin*_, which is discussed in Section 3.3. Fig. 2c illustrates the simulated *A*[*z*, Δ*j*_*r*_]_*noise*_*lin*_ after the same respective processing and analysis as Figs. 2a, without the LI of *e*[*j*]_*noise*_. Fig. 2d illustrates the simulated *A*[Δ*j*_*r*_]_*noise*_*lin*_ calculated from Fig. 2c and the predicted SDBG bias. It is clear that all the noises have the same amplitude, regardless of their center wavelengths. This matches the predicted SDBG bias since *e*[*j*]_*noise*_ is not correlated with *k* and no *k* distortion is present.

### 3.1. Experimentally measured SDBG

We applied the same analysis to experimentally acquired *e*[*j*]_*noise*_ using our vis-OCT system. Fig. 3a shows the *A*[*z*, Δ*j*_*r*_]_*noise*_*lin*_ from the measured *e*[*j*]_*noise*_, plotted using the same colorbar in Fig. 2a. The experimental *A*[*z*, Δ*j*_*r*_]_*noise*_*lin*_ in Fig. 3a has a similar, monotonic spectral bias to that of the simulated version (Fig. 2a). We confirmed this trend by measuring *A*[Δ*j*_*r*_]_*noise*_*lin*_ (Fig. 3b, red dashed line). Similar to the simulated *A*[Δ*j*_*r*_]_*noise*_*lin*_, the experimental *A*[Δ*j*_*r*_]_*noise*_*lin*_ (also for depths 500 *μm* − 800 *μm*) decreases approximately linearly with increasing central wavelength. Unlike the simulated *A*[*z*, Δ*j*_*r*_]_*noise*_*lin*_, the experimental *A*[*z*, Δ*j*_*r*_]_*noise*_*lin*_ (Fig. 3a) is not approximately constant with depth. Instead, its amplitudes are higher at shorter depths and decay exponentially with depth, which may be caused by two experimental conditions. First, the noise distribution from a supercontinuum laser is pink [22], which contains a higher proportion of low-frequency noises. Second, imperfect normalization of the light source spectral shape can propagate a low-frequency component into interference fringe. As shown in Eqn. 10, the SDBG bias is contingent on the spectroscopic processing of white noise. To this end, it was important to directly measure *A*[*z*, Δ*j*_*r*_]_*noise*_*lin*_ to monitor any inconsistencies with the model. Since the measured SDBG bias was in strong agreement with our mathematical model and simulation, we concluded, to a reasonable approximation, that background noise in our vis-OCT system was dominated by a white noise process. We did note small differences (~ 2% error) between the measured *A*[Δ*j*_*r*_]_*noise*_*lin*_ and predicted SDBG bias, which was likely explained by the lower noise frequencies, normalization of the light source shape, or *σ* not being perfectly constant across spectral locations. Since supercontinuum laser sources may vary in power, spectral shape, repetition rate, and RIN noise, it is important that researchers directly measure the SDBG.

**Fig. 3.**
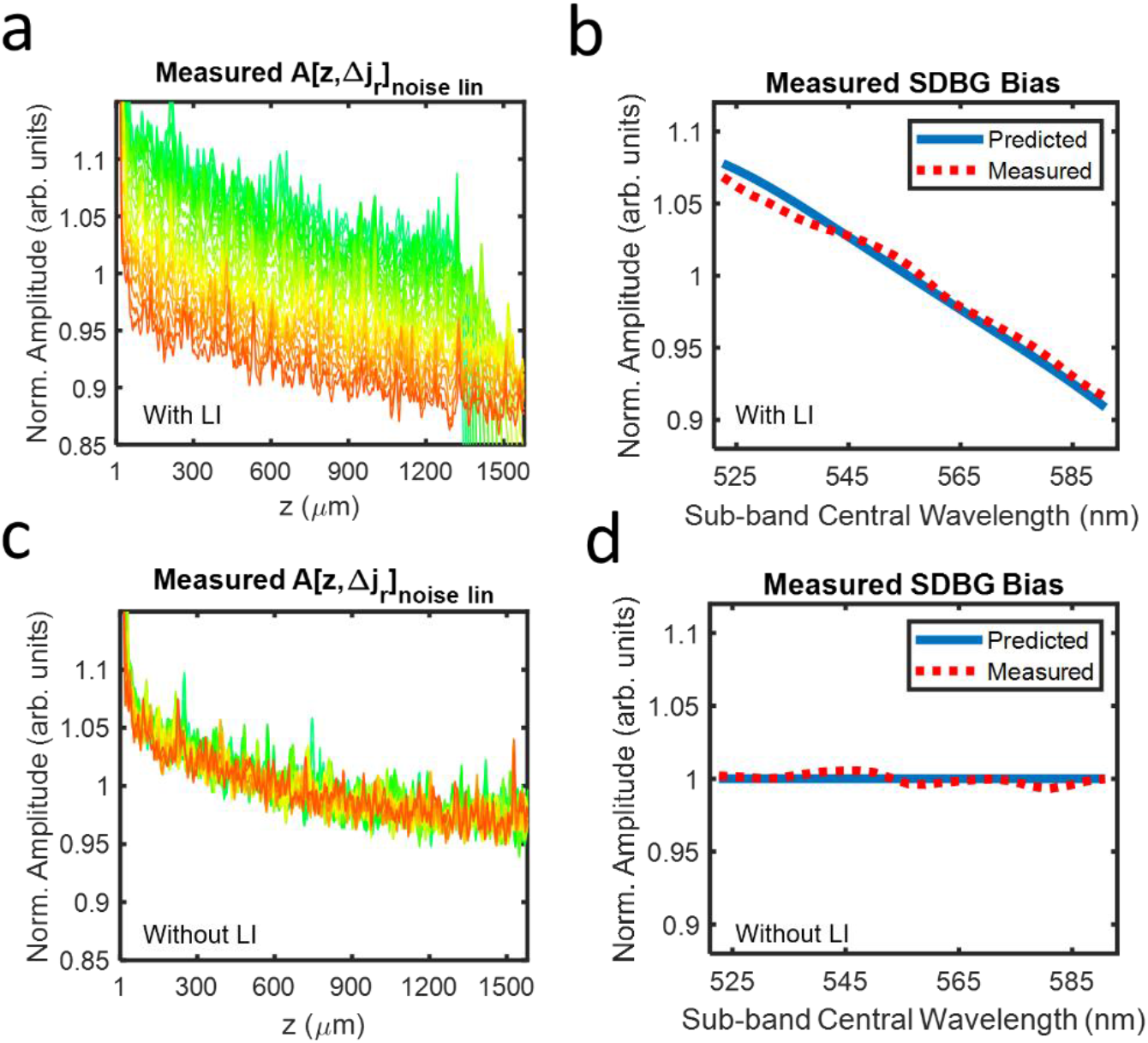
Measured SDBGs in vis-OCT. (a) Measured background SDA-lines with LI; (b) predicted SDBG bias (blue line) and measured SDBG bias (red dashed line) with LI from z = 500 μm to 800 μm; (c) measured background SDA-lines without LI; (d) predicted SDBG bias (blue line) and measured SDBG bias (red dashed line) without LI from z = 500 μm to 800 μm

Finally, Figs. 3c and 3d illustrate the measured *A*[*z*, Δ*j*_*r*_]_*noise*_*lin*_ and *A*[Δ*j*_*r*_]_*noise*_*lin*_, respectively, without LI. As predicted, the spectral dependence of the background noise floor is not present. Furthermore, depth decay of the background still existed, since the frequency distribution of the background noise is not influenced by interpolation.

### 3.3. Influence of interference fringe up-sampling

To this point, to simplify the calculation of SDBG, we did not extend the analysis to the filter in Eqn. 5. Indeed, the convolution of *t*[*j*] with *e*[*j*]_*noise*_ adds additional depth-dependence and spectral dependence to the SDBG. The STFT of *t*[*j*] for LI is

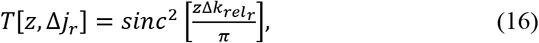

where *ξ* from Eqn. 5 is set to 1 since the total number of samples in each sub-band does not change. The multiplication of Δ*k*_*rel*_ with *z* signifies the scaling of the *k* domain after resampling. Therefore, true SDA-line can be written as

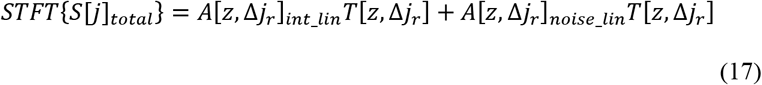

which adds the spectrally-dependent roll-off (SDR) [17] component to the signal and noise, according to Eqn. 16. It has been shown that uniformly up-sampling *S*[*j*]_*total*_ before the interpolation step reduces A-line amplitude decay with depth [19]. This is because up-sampling compresses SDA-lines, but not the depth-resolved interpolation filter. Compression of imaging depths relative to the interpolation filter exposes them to less decay by the *sinc*^2^ function than without compression (no up-sampling).

If *S*[*j*]_*total*_ is up-sampled at a high enough rate, the SDR induced by the interpolation filter becomes small enough to be negligible for most applications. This is also why there is no visually noticeable decay by the *sinc*^2^ function in Fig. 3. However, to our knowledge, an optimal up-sampling rate for spectroscopic vis-OCT has not been determined. An optimal up-sampling rate should be established since researchers may apply low upsampling rates to increase processing speeds without considering the spectroscopic consequences.

We investigate a minimum upsampling rate for spectroscopic vis-OCT in Fig. 4, which uses the simulation from Fig. 2a, except that we varied the up-sampling rates before interpolation. We compared *A*[*z*, Δ*j*_*r*_]_*noise*_*lin*_ *T*[*z*, Δ*j*_*r*_] corresponding to upsampling rates of 1, 2, 4, and 6. In Fig. 4a, *A*[*z*, Δ*j*_*r*_]_*noise*_*lin*_*T*[*z*, Δ*j*_*r*_] is influenced by the depth-independent amplitude scaling in Eqn. 13 and the depth-dependent interpolation filter in Eq 16. Each noise component decays with depth, where shorter wavelengths decay more rapidly than longer ones. Such decay adds two additional influences to the noise floor: decay with depth and different depth decay rates with different wavelengths. These influences further complicate the alterations of spectroscopic vis-OCT measurements, particularly when one tries to correct for them. They also complicate the SDBG correction strategy, which is described in Section 3.4. As shown in Figs. 4b and 4c, the influence of the interpolation filter is reduced with increased upsampling but cannot be completely removed. Under six-fold up-sampling (Fig. 4d), *A*[*z*, Δ*j*_*r*_]_*noise*_*lin*_*T*[*z*, Δ*j*_*r*_] only shows minute decay with depth, which suggests that *S*[*j*]_*total*_ should be up-sampled at least six times. At this point, Eqn. 17 can be simplified back to Eqn. 14. Finally, we note that *T*[*z*, Δ*j*_*r*_] is also multiplicative with the signal-carrying SDA-lines in Eqn. 17. Up-sampling *S*[*j*]_*total*_ by six-fold will reduce SDR of the signal-carrying SDA-lines in the same way as the background shown in Fig. 4.

**Fig. 4.**
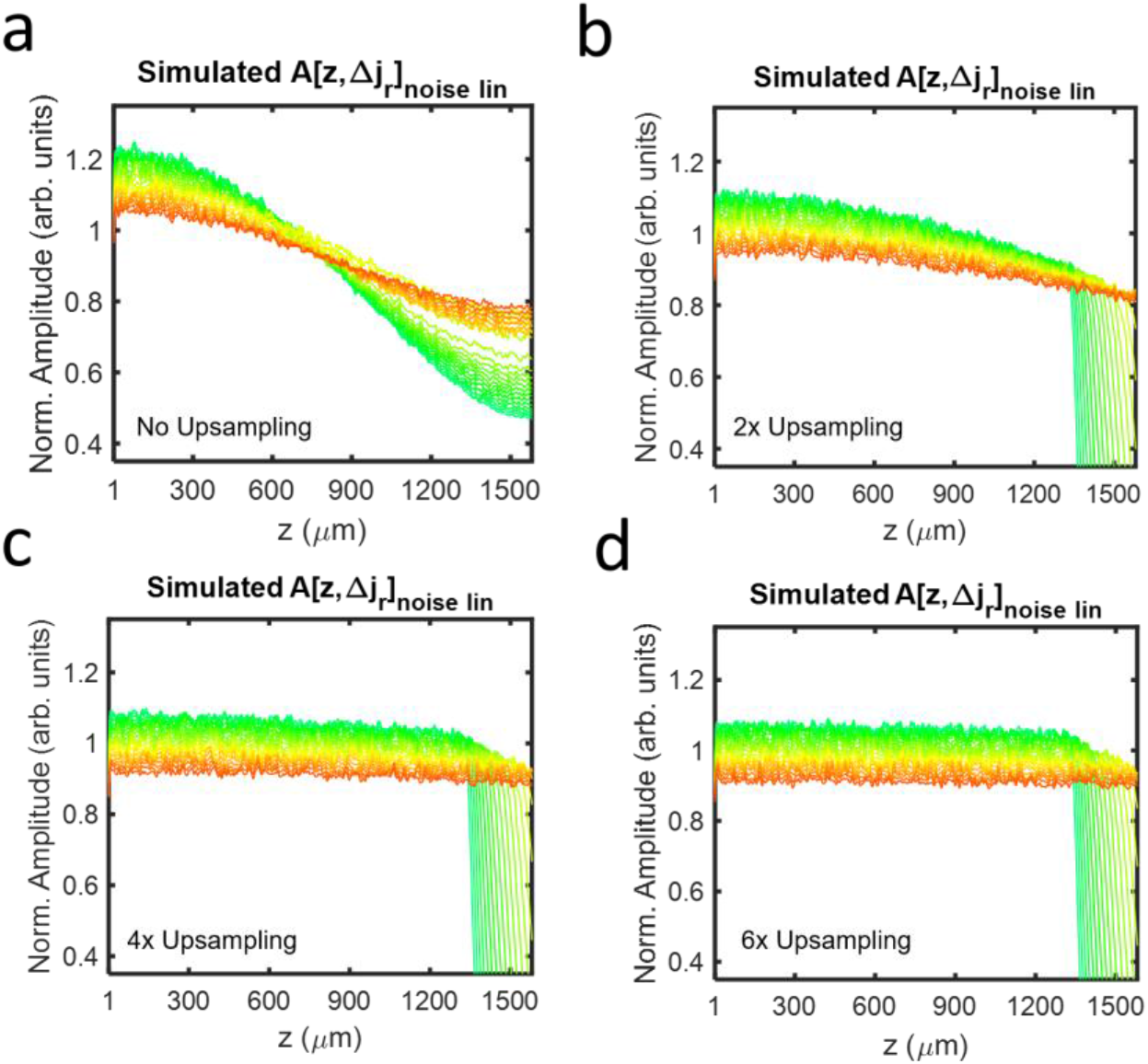
Simulated background SDA-lines with LI after (a) no, (b) two-fold, (c) four-fold, and (d) six-fold up-sampling.

### 3.4. Correcting SDBG in vis-OCT oximetry in humans

Influence of SDBG on spectroscopic analysis can be corrected experimentally by subtracting *A*[*z*, Δ*j*_*r*_]_*int*_*lin*_ from Eqn. 14 as

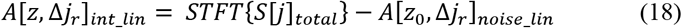

where *z*_0_ is the depth of the spectroscopic calculation. Practically, this can be accomplished by selecting a depth region where the structural vis-OCT signal is completely attenuated. Since the experimental SDBG was shown to decay with depth, we fit an exponential curve to each *A*[*z*, Δ*j*_*r*_]_*noise*_*lin*_ in the selected depth region. We then used the fitted values at *z*_0_ to estimate *A*[*z*_0_, Δ*j*_*r*_]_*noise*_*lin*_. We note that zero-padding the fringe at least 6-fold greatly simplifies this correction, since the depth decays of the SDBG are minimized and nearly uniform across all spectral sub-bands. When the vis-OCT signal is not completely attenuated, it is an acceptable alternative to acquire only the background signal, as shown in Fig. 3a, and directly calculate *A*[*z*_0_, Δ*j*_*r*_]_*noise*_*lin*_. However, it will also require knowledge of the DC sample arm spectrum, which is discussed in Section 4.

To demonstrate the importance of SDBG correction, we measured the vis-OCT spectrum of blood in human retinal vessels. Briefly, we imaged the retina of a healthy 23-year-old volunteer with a vis-OCT system described in [23]. Human imaging was approved by the Northwestern University Institutional Review Board and adhered to the Declaration of Helsinki. The optical power incident on the cornea was 250 *μW*. We measured the spectrum inside the retinal blood vessels using STFT sub-bands *r* = 3 to *r* = 23 (528 nm to 588 nm). We applied standard OCT processing, including removal of the spectrum DC component, 6-fold zero-padding, compensation for dispersion mismatch, and correction for the system roll-off. We averaged ~ 500 pixels in each vessel from 16 B-scans (8192 A-lines per B-scan) to reduce the background and speckle noise fluctuations.

We calculated the influence of SDBG in an artery (red box) and vein (blue box), as highlighted in Fig. 5a, near the optic disk. The spectrum was detected at an average distance of ~40 *μm* below the anterior vessel wall and is plotted in blue in Fig. 5b. All spectra are plotted after normalizing by their respective mean amplitudes. The red dashed-line in Fig. 5b is the predicted spectrum from the literature [24] for oxygen concentration (sO_2_) = 100%. Note how the blue spectrum is tilted downwards with increasing wavelength and does not well match the literature spectrum. Fig. 5c shows the measured *A*[*z*_0_, Δ*j*_*r*_]_*noise*_*lin*_. It matches well with those simulated and experimental SDBGs shown in Figs. 2b and 3b. The downwards tilt in the measured spectrum in Fig. 5b follows the trend of *A*[*z*_0_, Δ*j*_*r*_]_*noise*_*lin*_ in Fig. 5c.

**Figure 5.**
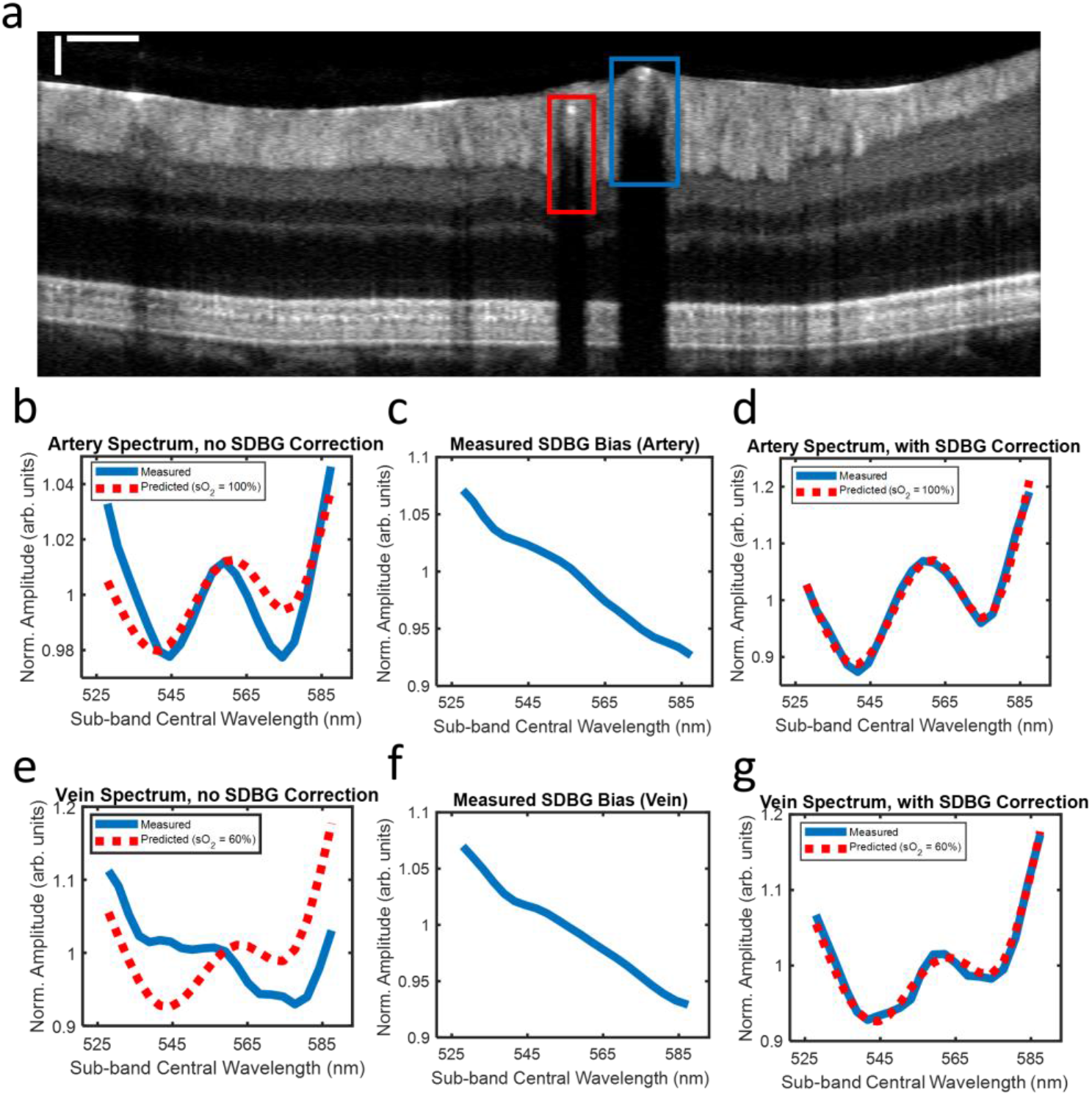
SDBG correction in the human retina. (a) B-scan image with one artery (red box) and one vein (blue box); scale bars are 250 *μm* (lateral) and 50 *μm* (axial); (b) measured blood spectrum in the artery before SDBG correction; (c) measured SDBG bias in the artery; (d) measured blood spectrum in the artery after SDBG correction; (e) measured blood spectrum in the vein before SDBG correction; (f) measured SDBG bias in the vein; (g) measured blood spectrum in the vein after SDBG correction

Fig. 5d shows the same spectral calculation in Fig. 5b after SDBG correction. We corrected the spectrum using *A*[*z*_0_, Δ*j*_*r*_]_*noise*_*lin*_ from Fig. 5c. After correction, the spectrum matches well (*R*^2^ = 0.98) with the literature spectrum (sO_2_ = 100%). Fig. 5e shows the measured spectrum (blue line) without SDBG correction in the vein and literature spectrum (red dashed-line) for sO_2_ = 60%. Again, the measured spectrum differs from the literature spectrum with a tilt downwards at longer wavelengths. Fig. 5f shows the measured *A*[*z*_0_, Δ*j*_*r*_]_*noise*_*lin*_. Fig. 5g shows the SDBG corrected spectrum, which well agrees with the literature spectrum (*R*^2^ = 0.98).

### 3.4. Metric to evaluate the influence of SDBG on spectroscopic OCT measurements

We recognized that the influence of SDBG also depends on the amplitude of the spectroscopic signal relative to the noise floor. To quantify SDBG’s influence on retinal oximetry in vessels similar to the ones in Fig. 5, we simulated oxygen-dependent spectra for sO_2_ = 0 to 100% and SDBGs for different signal-to-noise-floor-ratios (SNFRs). We define SNFR as

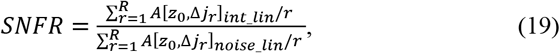

which is the ratio between the mean spectral amplitudes from the structure of interest and the noise floor. We calculated sO_2_ from the simulated spectra and calculated the root-mean-squared-error (RMSE) between the calculated sO_2_ and the ground truth. Figure 6 illustrates RMSE from the computer-generated ground truth when SNFR varies from 3 to 100 (sO_2_ values for SNFRs below 3 could not be quantified). Indeed, the error of sO_2_ is severely compromised by SDBG until SNFR > 55 (RMSE < 2%). For our vis-OCT setup, we found that typical SNFR in locations used for sO_2_ were no higher than 5, which was limited by the rapid attenuation of blood. The SNFRs in Fig. 5a were 2.05 and 1.77 for the artery and vein, respectively. The sO_2_ could not be calculated in these vessels until the SDBG was corrected. Therefore, we conclude that SDBG correction is critical for human retinal oximetry. The influence of SDBG correction will be different depending on specific applications. We recommend readers to simulate an accuracy metric similar to that shown in Fig. 6 to quantify the influence of the SDBG in a specific application.

**Figure 6.**
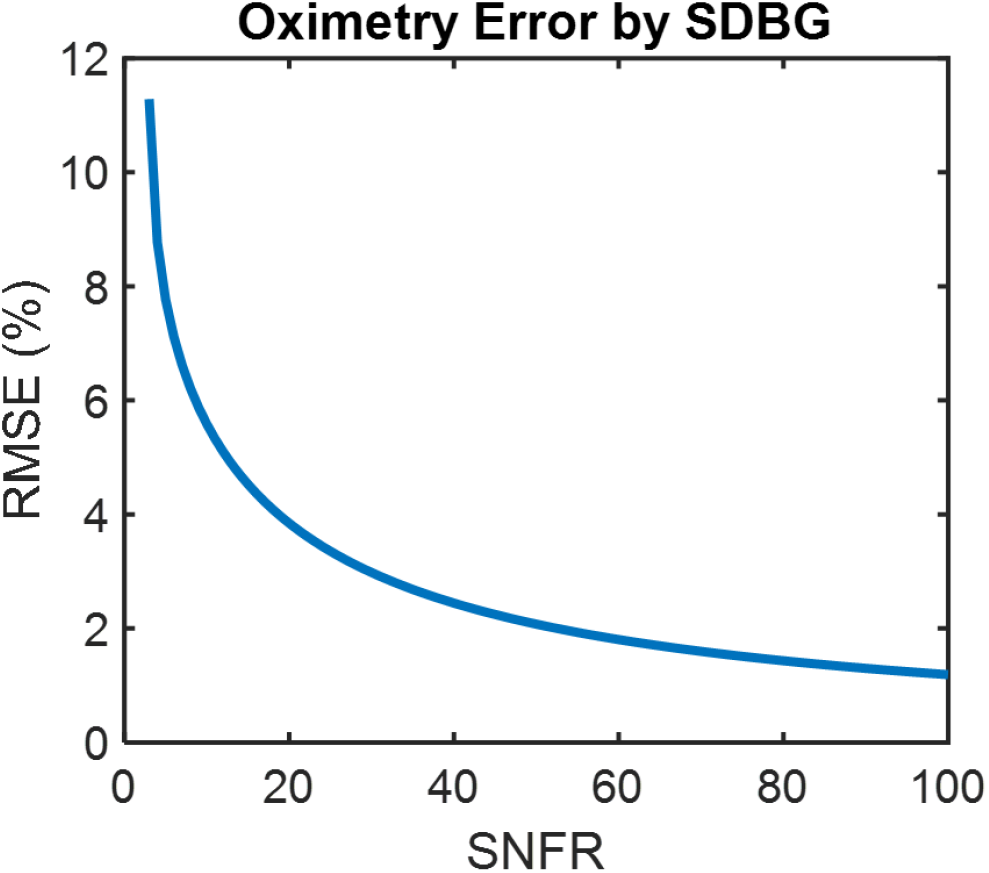
RMSE of simulated sO_2_ measurements with respect to the ground truth as a function of SNFR.

## 4. Discussion

We thoroughly investigated a systemic SDBG in spectroscopic SD-OCT. We developed a mathematical model to show that SDBG is caused by linear-in-*k* interpolation of the white noise. We validated the model using simulated and experimental SDBGs in vis-OCT. We demonstrated the importance of up-sampling the fringe before LI. We corrected SDBG in human spectroscopic vis-OCT and found that correcting SDBG was critical for measuring an accurate blood spectrum in humans. Finally, we showed the influence of SDBG in spectroscopic SD-OCT under different SNFR levels.

Potentially most insidious are cases where measured spectra agree with their literature models but are still subject to SDBG bias. Indeed, the SDBG amplitude derived here monotonically decreases with wavelength. Such shape is inconveniently correlated with the reported *λ*^−*α*^ scattering and backscattering spectra of many biological tissues, where *α* is a constant [3, 25]. As such, perceived scattering properties of tissue may be altered by the SDBG. This is relevant in a model of light-tissue interaction where the scattering coefficient or elements of the scattering coefficient a fitted parameter [12].

Finally, we note that *δk*_*rel*_ varies more in visible-light spectral range than in NIR spectral range. To this end, the SDBG is likely influence spectroscopic vis-OCT applications the most. Using a linear-in-*k* spectrometer [26] or swept-source [15] (not currently available in visible light), rather than a grating-based spectrometer, will greatly reduce the influence of SDBG since *δk*_*rel*_ becomes nearly constant. In this work, we performed all STFT analysis using sub-bands with uniform bandwidths in *k* space. Varying sub-bands’ bandwidths in the STFT to compensate for SDBG remains an open question. However, doing so will alter the amplitudes and resolutions of *A*[*z*, Δ*j*_*r*_]_*int*_*lin*_. Finally, we recognize that in spectroscopic OCT, the interference fringe is often divided by the source spectrum, 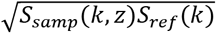 from Eqn. 1, for normalization purposes. As such, the measured SDBG may be altered by 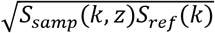, depending on normalization method and SDBG amplitude. However, such normalization does not alter the analysis concluded in this work, nor does it alter the proposed SDBG correction.

## 5. Conclusion

In conclusion, we demonstrated that spectroscopic SD-OCT signals are systemically biased by the spectrometer’s *k* spacing. We investigated this phenomenon in vis-OCT and found strong agreements between mathematical model, simulation, and experiment. We found that correction of the SDBG is important for retinal oximetry, a primary application for vis-OCT. By establishing and verifying the principles of the SDBG, this work informs researchers towards making accurate spectroscopic OCT measurements.

## Acknowledgments

The authors thank David A. Miller and Lisa Beckmann for helpful discussions.

